# N-glycans show distinct spatial distribution in mouse brain

**DOI:** 10.1101/2023.05.30.542954

**Authors:** Maxence Noel, Richard D. Cummings, Robert G. Mealer

## Abstract

Protein N-linked glycosylation is a ubiquitous modification in the secretory pathway that plays a critical role in the development and function of the brain. N-glycans have a distinct composition and undergo tight regulation in the brain, but the spatial distribution of these structures remains relatively unexplored. Here, we systematically employed carbohydrate binding lectins with differing specificities to various classes of N-glycans and appropriate controls to identify multiple regions of the mouse brain. Lectins binding high-mannose-type N-glycans, the most abundant class of brain N-glycans, showed diffuse staining with some punctate structures observed on high magnification. Lectins binding specific motifs of complex N-glycans, including fucose and bisecting GlcNAc, showed more partitioned labeling, including to the synapse-rich molecular layer of the cerebellum. Understanding the distribution of N-glycans across the brain will aid future studies of these critical protein modifications in development and disease of the brain.

## INTRODUCTION

Protein glycosylation is essential in the brain. Severe impairments of the pathway caused by Mendelian disorders of glycosylation lead to profound neurodevelopmental abnormalities^1^ and a growing body of evidence supports its role in more complex neuropsychiatric phenotypes ranging from prion disorders^2^ to Alzheimer’s disease^3^ and schizophrenia.^4^ Asparagine (N-) linked protein glycosylation has been a major focus of study in the brain^5^ facilitated by the ubiquitous nature of the modification, a well-defined synthetic pathway, and the facile use of tools including Peptide:N-glycosidase F (PNGase F) to release N-glycans from the protein backbone for analysis. Most work has focused on single glycan modifications in isolation or surveys on the pool of glycans present en masse. However, such studies often require molecular tools present only in laboratories focused on glycobiology and a niche understanding of the proper experimental controls for carbohydrate chemistry.

Lectins, carbohydrate binding proteins often isolated from plants, are powerful tools in glycobiology due their relatively specific affinities to diverse types of structures.^6^ However, lectins often possess a range of affinity for numerous glycans that may or may not have structural or compositional similarity, which can lead to a misinterpretation of their signal in novel contexts such as different species, tissues, and assays.^7^ Lectin microarrays have been useful in examining the bulk glycan profiles across tissues.^8^ Pandit and colleagues presented a lectin fluorescence protocol for use in tissue sections including brain, importantly emphasizing positive and negative controls, though primary brain data was not included beyond a single representative image.^9^ Their related study investigating N-glycans in the striatum and substantia nigra nicely correlated LC-MS data and fluorescence imaging for three lectins, but did not include controls for binding, different magnification levels, or additional brain regions.^10^

In this study, we combined a panel of frequently used N-glycan binding lectins with histochemical labeling of fixed mouse brain tissue. Following optimization and confirming specificity of each lectin, detailed imaging analysis was performed across the brain with the lectins abbreviated ConA, GNL, AAL, PHA-E, RCA, SNA, and MAL-I. Descriptions of lectin binding within the cerebellar layers are included, as well as general patterns of binding across the brain and several structures of note. These findings highlight the unique and non-uniform spatial distribution of N-glycans across the brain and provide an adaptable tool that can be utilized for the study of N-glycans in neuroscience research.

## EXPERIMENTAL METHODS

### Mouse samples

All mice were maintained in standard housing conditions, provided normal chow and water *ad libitum*, and maintained on 12-hour day/light cycle. All mice were housed and maintained in accordance with the guidelines established by the Animal Care and Use Committee at Beth Israel Deaconess Medical Center under the approved protocol #02-2022. Twelve-week-old male mice were used in this study unless otherwise indicated.

### Histology/fluorescence imaging

Mice at 12 weeks of age were euthanized with CO_2_ gas in accordance with AVMA guidelines and trans-cardially perfused with ice-cold PBS containing heparin at 10,000 units/L (Sigma #3149) for 2 minutes at 6 ml/minute, followed by ice-cold 4% PFA in 0.1 M PBS, pH 7.4 for 5 minutes at 5 ml/minute. Whole brains were then removed from the skull and postfixed in 4% PFA overnight and transferred to PBS until processing. Fixed brain was paraffin embedded and cut in 3 μm coronal sections with a microtome at the BIDMC Pathology Core Facility. Slides were then deparaffinized using a standard xylene/ethanol gradient protocol and placed in antigen retrieval solution (0.1 M citric acid and 0.1 M sodium citrate, pH=6), incubated in pressure cooker at boiling temperature for 3 min, and stored at 4°C in TBS. Tissue sections were circled by hydrophobic PAP PEN and slides were treated with denaturing buffer for 5 min at 95°C. Sections were washed with TBS (Trizma 20 mM, NaCl 100 mM, CaCl_2_ 1 mM, MgCl_2_ 1 mM, pH 7.2) 3 times followed by 3 washes with TBS-tween 0.05% (TBS-T). Sections were blocked with 3% BSA in TBS for 1 h at room temperature (RT). Tissues were washed again three times with TBS and incubated with 25 μg/mL of lectins (Vector Labs: ConA FL-1001, GNL B-1245, AAL FL-1391, PHA-E FL-1121, RCA-I B-1085, SNA FL-1301, MAL-I FL-1311) in TBS-T for 1 h at RT in darkness with gentle shaking. For control glycosidase experiments, slides were pretreated with 2,500 U of PNGase F in a final volume of 80 μL at 37°C for 1 hour prior to incubation with lectins. For control carbohydrate/haptenic sugar blocking experiments, lectins were incubated for 30 min with the indicated sugar before incubation on the slide for 1 hour at RT. After the incubation with lectins, sections were washed with TBS-T 3 times. Most lectins described above were directly conjugated to FITC – however, for biotinylated RCA-I and GNL, streptavidin-488 (Invitrogen, S11223) was used as secondary antibody at 10 μg/mL and incubated in TBS-T for 1 hour at 25°C prior to imaging. Sections were washed 2 times with TBS-T, one time with TBS and counterstained with Hoechst 33342 diluted at 1/1000 in TBS for 10 min at RT in darkness. Sections were then washed 3 times with TBS, incubated with 0.1% Sudan Black B (Sigma, 199664) in 70% ethanol for 5 min to reduce autofluorescence coming from lipofuscin, washed again 2 times with TBS and mounted with a glass coverslip using Prolong Gold Antifade Mountant (ThermoFisher, P36930). After slides cured overnight, image acquisition was performed on Zeiss LSM 880 Up-right Confocal System at the BIDMC Confocal Microscopy Core for high magnification and a VS120 Slidescanner from Olympus for low magnification. Images analysis was performed using ImageJ (v1.53t) or QuPath (v0.3.2) software.

## RESULTS

To ensure correct interpretation of lectin staining to coronal mouse brain sections, we generated a standardized protocol which was then optimized for each lectin on sections deprived of glycolipids after tissue processing. After maximizing binding affinity, specificity was confirmed by both sensitivity to glycosidase treatment and inhibition with competing carbohydrates (haptens) based on previous characterizations. In addition, fluorescently labeled and biotinylated lectins, which are commercially available, were confirmed for their specificity by glycan microarray analyses using resources at the National Center for Functional Glycomics (NCFG) (Fig. 1A). The lectins used and references for their specificities include Concanavalin A (ConA),^11,12^ *Galanthus nivalis* lectin (GNL),^13,14^ *Aleuria aurantia* lectin (AAL),^15^ *Phaseolus vulgaris* erythroagglutinin-E (PHA-E),^16^ *Ricinus communis* agglutinin-I (RCA),^11^ *Sambucus nigra* agglutinin (SNA),^17^ and *Maackia amurensis* lectin (MAL-I).^18,19^ Using cerebellar tissue, the binding of the commonly used lectin ConA to the mannose core of most N-glycans was confirmed by signal loss following treatment with Peptide:N-glycosidase F (PNGase F) that removes all types of N-glycans, and by competitive inhibition of lectin binding with 200 mM glucose and 200 mM mannose (Fig. 1B).

**Figure 1.**
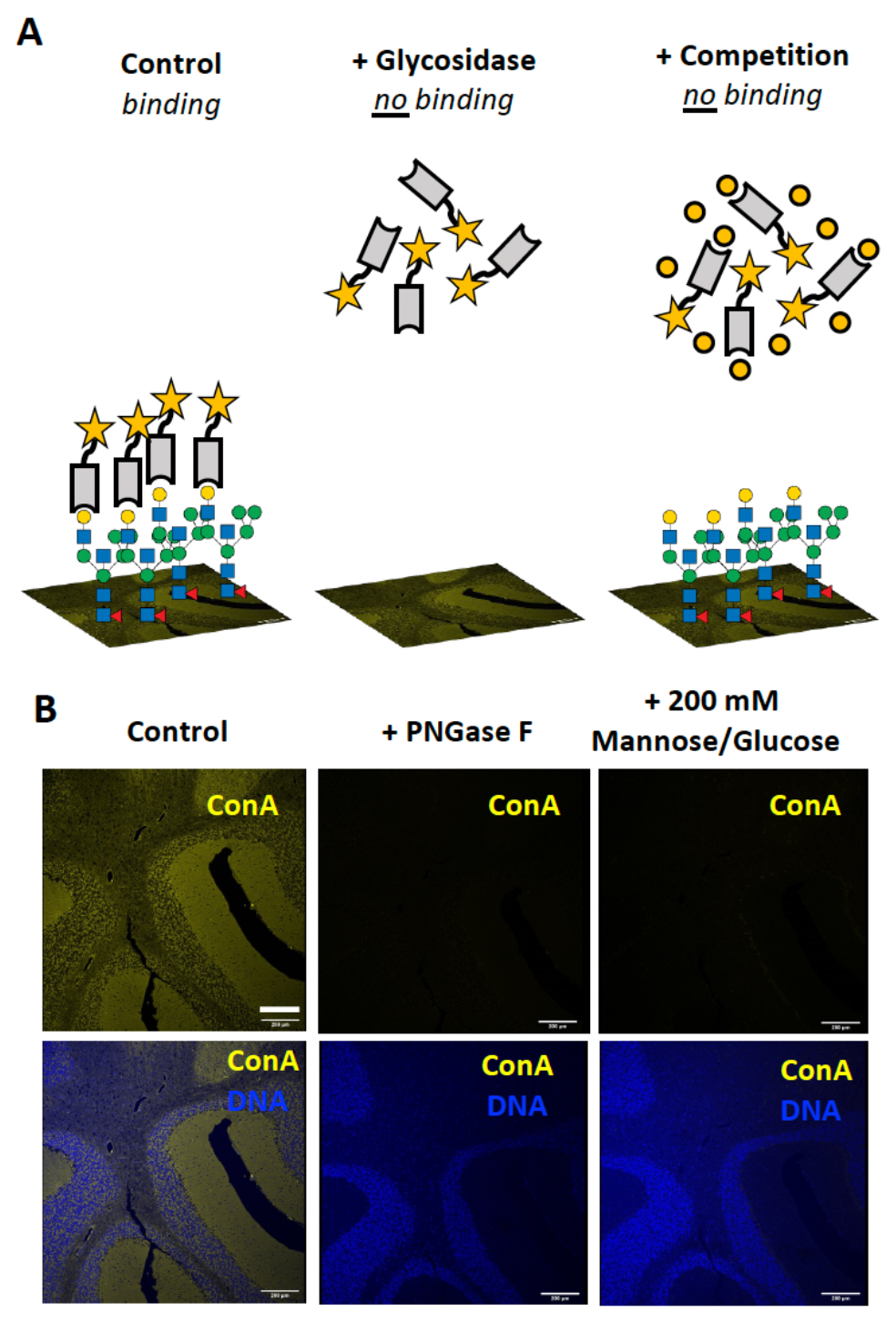
Optimized lectin fluorescence in brain slices. A) Schematic highlighting controls for determining specific binding of lectins to brain slices. The binding of lectins (gray) with a reporter such as a fluorescent tag (star) reveals the spatial distribution of glycan classes on a tissue slide. Specific binding of the lectin to the glycan structures is confirmed by determining its sensitivity to appropriate glycosidases as well as through inhibition under saturating concentrations of a competitive carbohydrate. B) The specificity of ConA binding to its substrate on the layers of the cerebellum is confirmed via removal of the N-glycans by incubation with PNGase F and inhibition of the lectin binding by saturating conditions of mannose and glucose (200 mM each). Scale bar = 200 μm.

High magnification images at the Purkinje cell layer junction of the cerebellum highlight several unique characteristics of N-glycan lectin binding patterns, including the presence of punctate structures, varying degrees of staining across anatomical layers, and binding to luminal structures (Fig. 2). We next focused on binding patterns across the brain, utilizing three coronal planes to survey numerous macroscopic structures including cerebellum, brain stem, corpus callosum, hippocampus, thalamus, hypothalamus, cortex, and olfactory bulb (Fig. 3A and 3B, Supp. Fig. 1). Differential binding patterns were observed for each lectin, as detailed below, and summarized in Table 1.

**Table 1.** N-glycan binding lectin properties in the brain. ^*^Binding pattern in coronal sections according to Supp. Fig. 1. Abbreviations: PNGase F - Peptide:N-glycosidase F; ML - molecular layer; GL - granular layer; AV- arbor vitae; WMT - white matter tracks.

| Lectin | PNGase F sensitivity | Competing sugar (200mM) | Regional Binding Pattern* |  |  | Other notable structures |
| --- | --- | --- | --- | --- | --- | --- |
|  |  |  | i | ii | iii |  |
| ConA | + | Mannose, Glucose | Diffuse in ML/GL, less in AV | Diffuse, less in WMT | Diffuse | Bright punctate structures, including around Purkinje cell body |
| GNL | + | Mannose, Glucose | Diffuse in ML/GL, less in AV | Diffuse, less in WMT | Diffuse with increased signal in glomerular layer | Bright punctate structures, including around Purkinje cell body |
| AAL | + | L-Fucose | Increased in ML, scattered in GL, decreased in AV | Diffuse, less in WMT | Diffuse | - |
| PHA-E | + | Lactose | Increased in ML, scattered in GL, decreased in AV | Diffuse | Diffuse | Bright signal in choroid plexus and moderate signal in microvasculature |
| RCA-I | + | Lactose, Galactose | Speckled throughout | Speckled throughout | Speckled with moderate signal in glomerular layer | Bright signal in choroid plexus and microvasculature |
| SNA | Partial | Lactose | Specific binding in ML and AV, non-specific in GL | Diffuse | Diffuse | Bright signal in choroid plexus |
| MAL-I | + | Lactose | Increased in ML, scattered in GL, decreased in AV | Diffuse, increased signal near surface | Diffuse, increased signal near surface | - |

**Figure 2.**
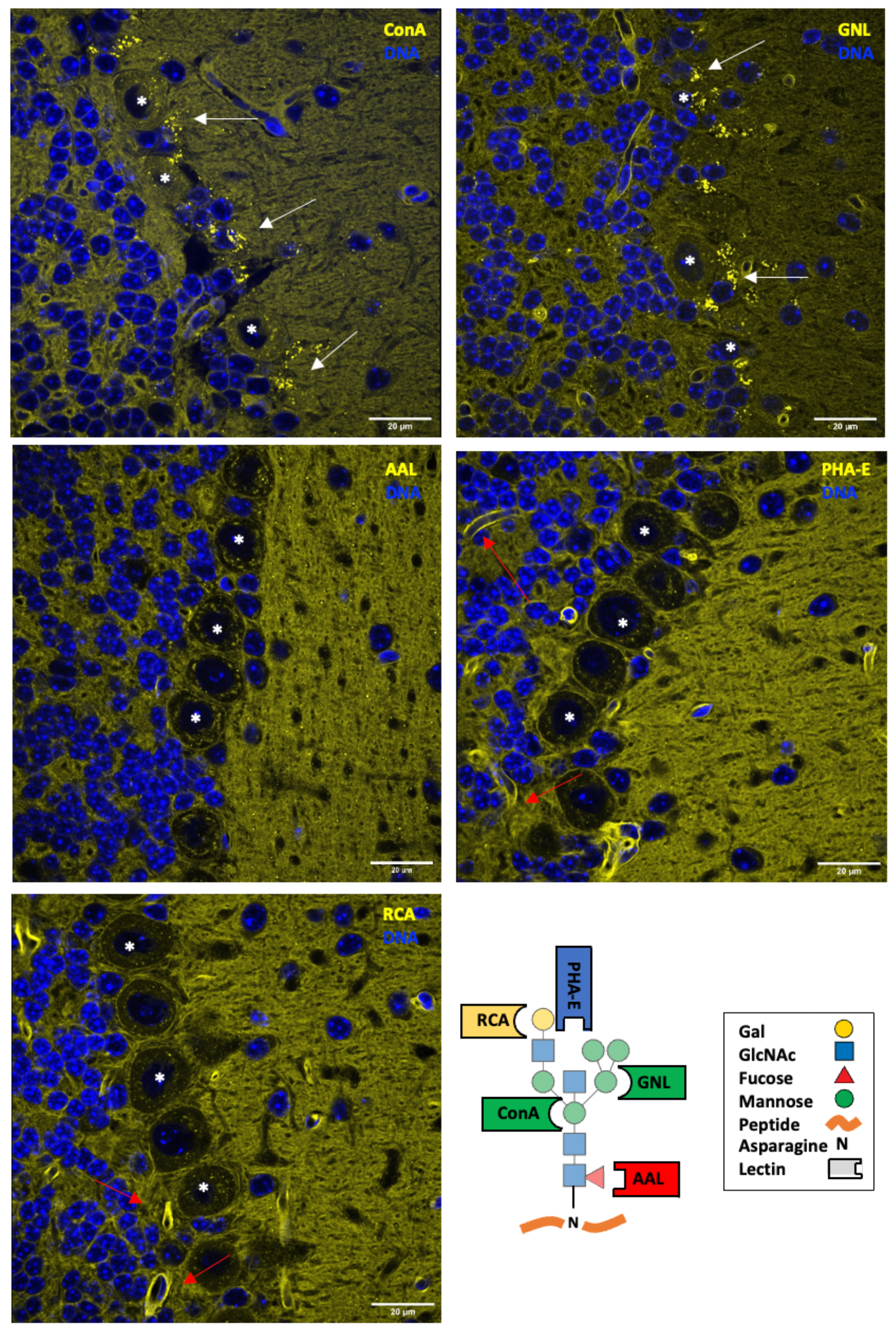
N-glycan binding lectins highlight distinct structural elements at the Purkinje cell junction of the cerebellum. In addition to diffuse binding across layers of the cerebellum, the lectins GNL and ConA showed bright punctate staining consistent with synapses (white arrows) around Purkinje cell bodies (^*^). Binding of AAL and PHA-E showed increased signal in the molecular layer with a few diffuse punctate structures noted. PHA-E and RCA showed bright signal in multiple luminal structures consistent with microvasculature (red arrows). Scale bar = 20 μm. A schematic with common lectin-binding sites is shown for reference.

**Figure 3.**
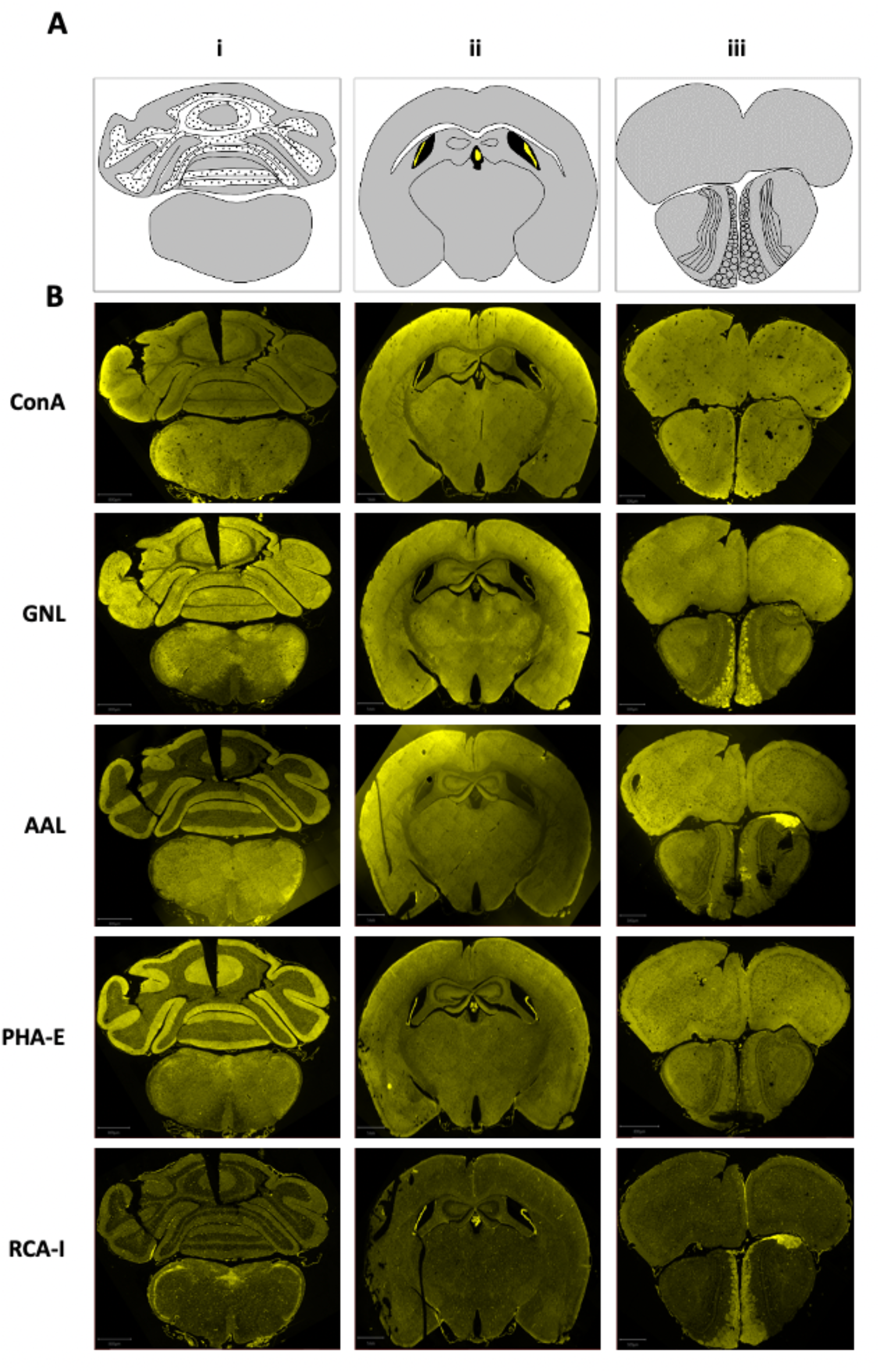
N-glycan binding lectins display distinctive patterns across the brain. A) Schematic of the different coronal brain sections, highlighting prominent structures including gray matter (gray), white matter (white), ventricles (black), and choroid (yellow). B) Staining of the different brain sections with the indicated lectins. Scale bars: i = 800 μm; ii = 1 mm, iii = 500 μm.

The lectin ConA, which binds mannose within high mannose, hybrid, and biantennary complex-type N-glycans, showed diffuse staining across nearly every region of the brain (Supp. Fig. 2). Specificity of the signal was confirmed by PNGase F treatment and inhibition with 200 mM mannose and 200 mM glucose. In the cerebellum, ConA binding appeared highest in the molecular layer (ML), with less in the granular layer (GL), and the least in the arbor vitae AV. This diffuse pattern was present across the more rostral brain structures, with a decreased signal present in white matter tracts including the corpus callosum. Several punctate structures were noted in and around the cell bodies of the Purkinje cell layer.

GNL, which binds high mannose and hybrid-type N-glycans, also showed diffuse distribution across the brain (Supp. Fig. 3). Specificity of the signal was confirmed by PNGase F treatment and inhibition with 200 mM mannose and 200 mM glucose, which removed the bulk of the signal. The minor residual background could result from incomplete digestion on the slide as this lectin binds the most abundant class of N-glycans in the brain, or simply represents residual non-specific signal. In the cerebellum, near equal intensity of GNL binding was observed in the ML and GL, with reduced intensity in the AV. Similar to ConA, high magnification showed punctate structures binding GNL in the Purkinje cell layer.

AAL, which binds nearly all linkages of fucose to N-glycans, showed broad staining but considerable layering in certain areas (Supp. Fig. 4). Specificity of the signal was confirmed by PNGase F treatment and inhibition with 200 mM L-fucose. In the cerebellum, AAL signal was dramatically enriched in the ML compared to both the GL and AV. Though some punctate structures were noted on high magnification, these appeared more diffusely spread across the ML.

PHA-E, which has affinity for bisected N-glycans containing a β-1,4 linked GlcNAc to the core mannose, showed a pattern similar to AAL (Supp. Fig. 5). Specificity of the signal was confirmed by PNGase F treatment and inhibition with 200 mM lactose. Intense binding was observed in the ML, with far less signal in both the GL and AV. Upon higher magnification in the cerebellum, several luminal structures were present suggesting some microvasculature was labeled by PHA-E, though this was not the bulk of the signal.

RCA, which has affinity to terminal β-linked galactose of N-glycans, showed a distinctive speckled pattern compared to the other lectins (Supp. Fig. 6). Specificity of the signal was confirmed by PNGase F treatment and inhibition with 200 mM lactose and 200 mM galactose. The speckled pattern, which was present across each region and contributed to most of the signal, was prominent in the cerebellum as luminal structures consistent with microvasculature. Minor signal was noted in the AV and ML with the least observed in the GL.

SNA, which is commonly used as a marker for glycans containing α-2,6 linked sialic acid,^7^ but can also bind sulfated N-glycans, showed relatively broad binding (Supp. Fig. 7). However, the specificity of the signal was not uniform, as PNGase F treatment and inhibition with 200 mM lactose failed to remove all binding. Strong SNA signal remained in the GL, while that in the ML and AV was reduced.

MAL-I, which binds α-2,3 sialic acid and sulfated glycans,^18^ exhibited a unique binding profile unique from the other lectins tested (Supp. Fig. 8). Binding specificity was confirmed with PNGase F and 200 mM lactose inhibition, with the latter condition leaving trace staining within Purkinje cell bodies. The most intense MAL-I binding was observed in the ML, with far less signal seen in other layers of the cerebellum and brain stem. On high magnification, several luminal structures were noted suggestive of some staining to the microvasculature.

## DISCUSSION

In prior studies of the brain protein N-glycome, measured with both MALDI-TOF MS and lectin blotting, we observed an abundance of N-glycans that comprised of high mannose, bisected, and fucosylated structures, with a scarcity of galactosylated and sialylated species.^20^ Here, we extend these findings to address the distribution of the major N-glycan classes using optimized lectin fluorescence across the mammalian brain. The patterns suggest a level of a spatial regulation related to their functions, with the bulk of N-glycans showing diffuse distribution and more complex and modified structures showing a more restricted pattern.

ConA binds most N-glycans and GNL binds the high mannose structures that comprise the bulk of brain N-glycans (∼60%); thus, their similar patterns are consistent with prior studies.^20^ The ConA+ and GNL+ puncta nearby Purkinje cell bodies suggest that these high mannose structures can be enriched in specific structures surrounding neuronal cell bodies such as synapses. AAL and PHA-E, which respectively show affinity to the fucosylated and bisected structures that make up the next major independent category of brain N-glycans (∼30%), also showed a complementary pattern of binding, supporting our previous findings of their shared presence on the same N-glycans.^20^ AAL and PHA-E also showed enrichment in the synapse rich molecular layer of the cerebellum. Bradberry *et al*. recently demonstrated the enrichment of fucosylated species within synaptosome structures analyzed by LC-MS.^21^ Though no mention was made of bisected structures, likely due to the challenge of determining the precise location of each additional GlcNAc residue, these findings appear consistent with our results of terminally modified N-glycans showing enrichment in synapse dense areas.

Localization of less abundant N-glycan classes, such as those containing galactose and sialic acid, present additional challenges for study. The strongest RCA signal was within micro-vasculature structures of the brain, in contrast with the other N-glycan binding lectins having high signal in the parenchyma itself. However, this result may be complicated by the presence of the abundant α-1,3 linked fucose to the galactose-GlcNAc-fucose containing Lewis X motif in brain, which likely impairs RCA binding based on data from the NCFG glycan array. RCA binding was enriched in some unique areas, such as the area postrema, a brain stem area of increased permeability, and ventricular choroid structures. The binding of SNA and MAL-I to sialic acid-containing glycans is challenging to interpret due to several factors. These N-glycans are of even lower abundance in the brain, and these lectins can have affinity to other structures such as sulfated glycans. However, both SNA and MAL-I showed affinity to ventricular choroid structures, and the striking binding of MAL-I almost exclusively to the molecular layer of the cerebellum is of particular interest and will warrant future studies. Minami *et al*., suggest that dynamic regulation of sialic acid is involved in memory processing, and show a decrease of MAL-I binding following acute hippocampal slice depolarization, though lectin controls were not presented.^22^

In summary, we present an N-glycan landscape across the murine brain, showing distinct structural and spatial distribution of N-glycan classes. These findings complement our studies and those of others using mass-spectrometry and lectin blotting to explore the unique characteristics of protein glycosylation in the mammalian brain,^20,23^ adding a validated and simple approach to explore glycan location. MALDI imaging is a promising and emerging tool for glycan visualization in fixed tissue that combines glycan release with mass spectrometry, generating a dataset with molecular resolution down to individual glycan masses.^24^ However, the spatial resolution of MALDI imaging is currently limited at best to the micrometer range, which prevents its application to specialized structures such as dendritic spines. Work by Hanus *et al*. with primary neurons highlights the intricacy of studying N-glycosylation in brain, as lectins showed differing degrees of synaptic and surface co-localization resulting from the unique secretory processing within these cultured cells.^25^ A full understanding of N-glycosylation in the brain will require a combination of techniques and models, each adding an essential piece to this complex puzzle.

## ASSOCIATED CONTENT

### Supporting Information

*Supplementary Figures S1*–*S8*. (PDF)

## AUTHOR INFORMATION

### Author Contributions

The manuscript was written through contributions of all authors. All authors have given approval to the final version of the manuscript.

### Notes

The authors declare no competing financial interest.

## ACKNOWLEDGMENT

We acknowledge the Protein-Glycan Interaction Resource of the CFG and the National Center for Functional Glycomics (NCFG) at Beth Israel Deaconess Medical Center, Harvard Medical School (supporting grant R24GM137763). We would like to that the Beth Israel Deaconess Medical Center Microscopy Core for technical support. RGM is supported by 1K08MH128712 and a 2021 NARSAD Young Investigator Grant from the Brain & Behavior Research Foundation. This research was supported by NIH Grant R24GM137763, awarded to RDC.

