## Supplementary Figures for "N-glycans show distinct spatial distribution in mouse brain"

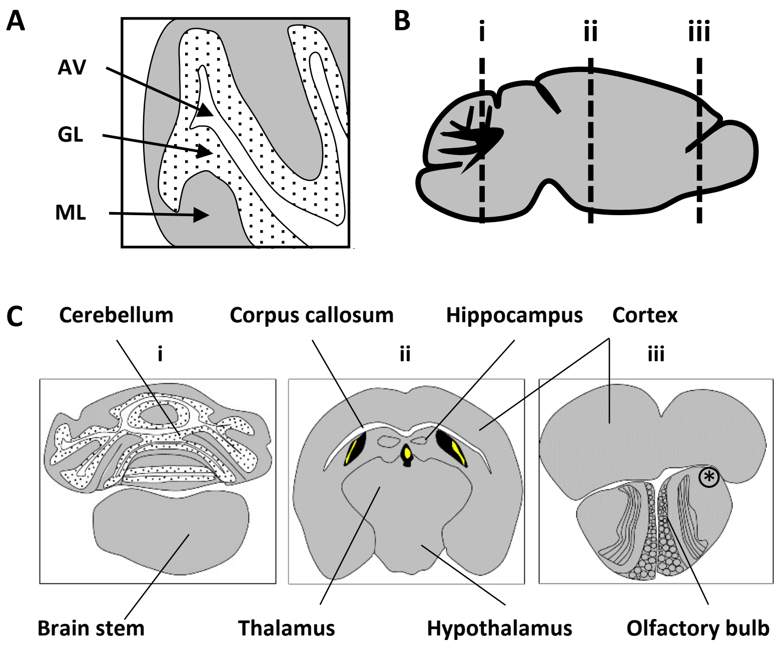


**Supplementary Figure 1 – Schematic Illustrations. A)** Layers of the cerebellum including the arbor vitae (AV), granular layer (GL), and molecular layer (ML). **B)** Approximate locations of coronal sections used for lectin staining. **C)** Annotation of the major brain structures across coronal brain sections noted in (B). Gray matter (gray), white matter (white), ventricles (black), and choroid (yellow) are noted, in addition to large regions including the cerebellum, brain stem, corpus callosum, hippocampus, thalamus, hypothalamus, cortex, and olfactory bulb are indicated. Of note, a unilateral, asymmetric bulge is noted above one of the olfactory bulbs (*), with staining suggesting this likely represents a blood clot/bleed that formed during perfusion.

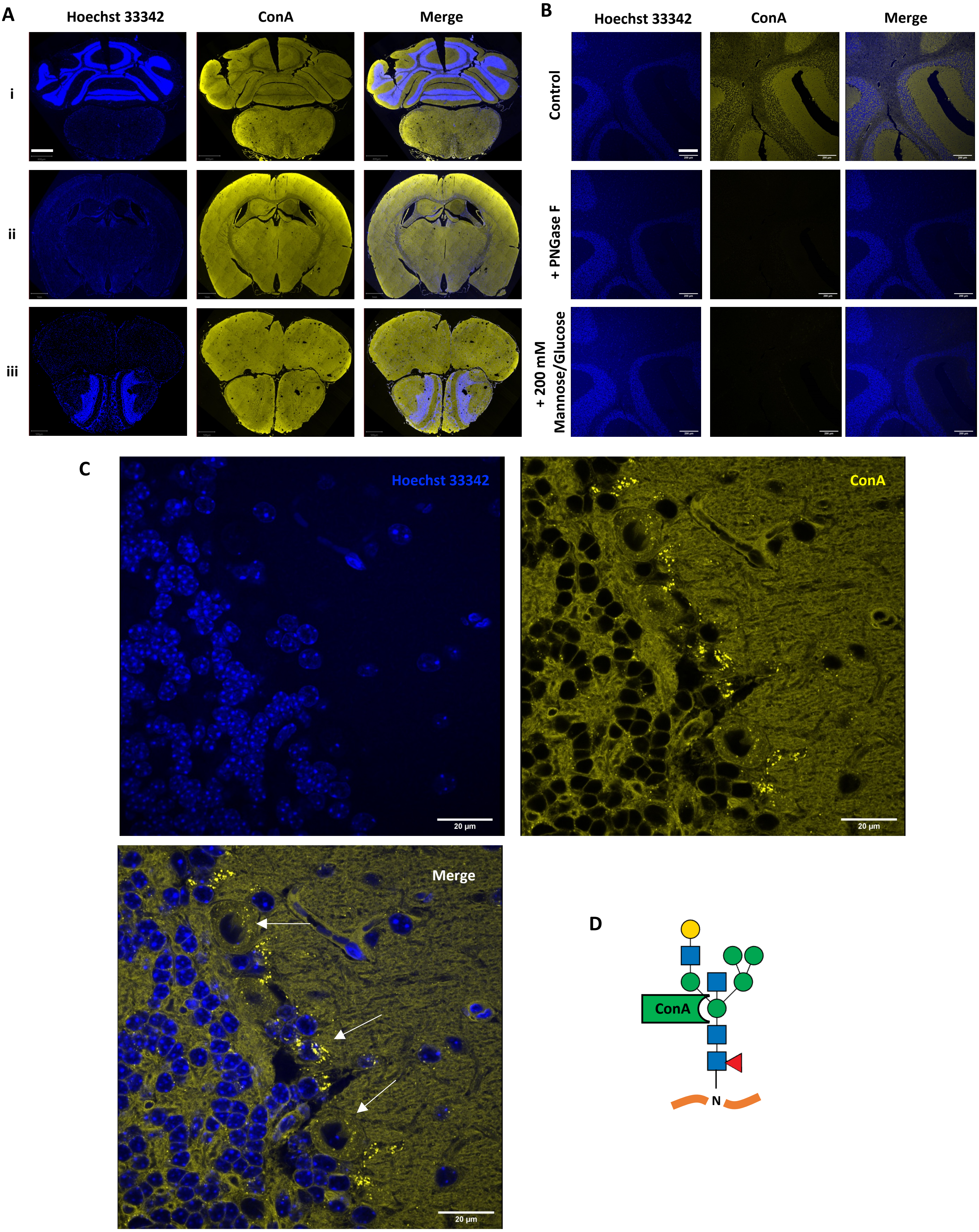


**Supplementary Figure 2 - ConA binding in the brain is broad and diffuse. A)** ConA binding across multiple brain regions corresponding to coronal sections (i, ii, iii) indicated in Supp. Fig. 1. Scale bars i = 800 µm, ii = 1 mm, iii = 500 µm. **B)** ConA binding specificity in the cerebellum is confirmed by sensitivity to PNGase F and competition with mannose and glucose. Scale bar = 200 µm. **C)** ConA binding of the Purkinje cell layer, highlighting punctate structures (white arrows) around several cell bodies. Scale bar = 20 µm. **D)** Binding preference of ConA to the core structure of N-glycans.

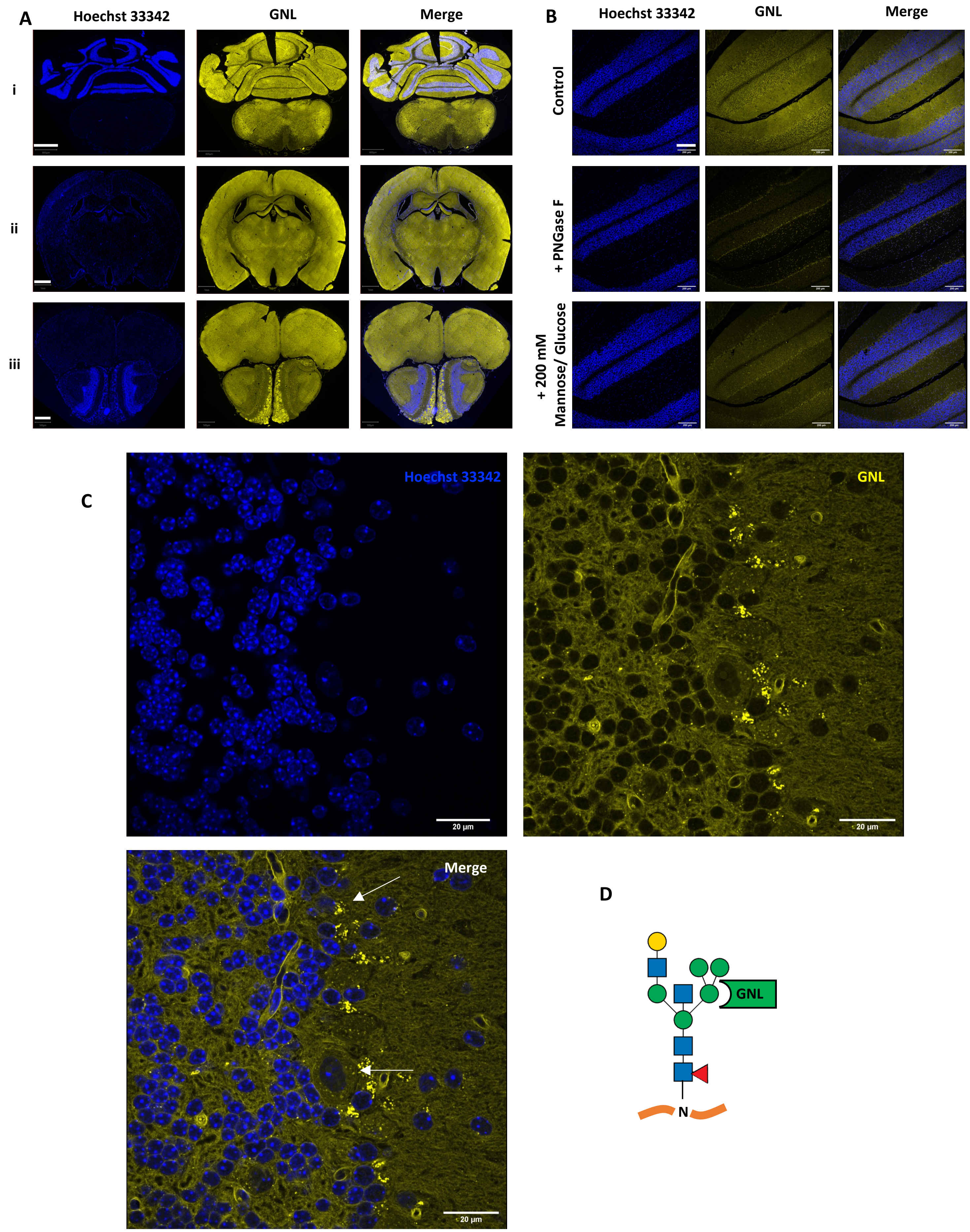


**Supplementary Figure 3 - GNL binding in the brain is broad and diffuse. A)** GNL binding across multiple brain regions corresponding to coronal sections (i, ii, iii) indicated in Supp. Fig. 1. Scale bars i = 800 µm, ii = 1 mm, iii = 500 µm. **B)** GNL binding specificity in the cerebellum is confirmed by sensitivity to PNGase F and competition with mannose and glucose. Scale bar = 200 µm. **C)** GNL binding of the Purkinje cell layer, highlighting punctate structures (white arrows) around several cell bodies. Scale bar = 20 µm. **D)** Binding preference of GNL to high-mannose type N-glycans.

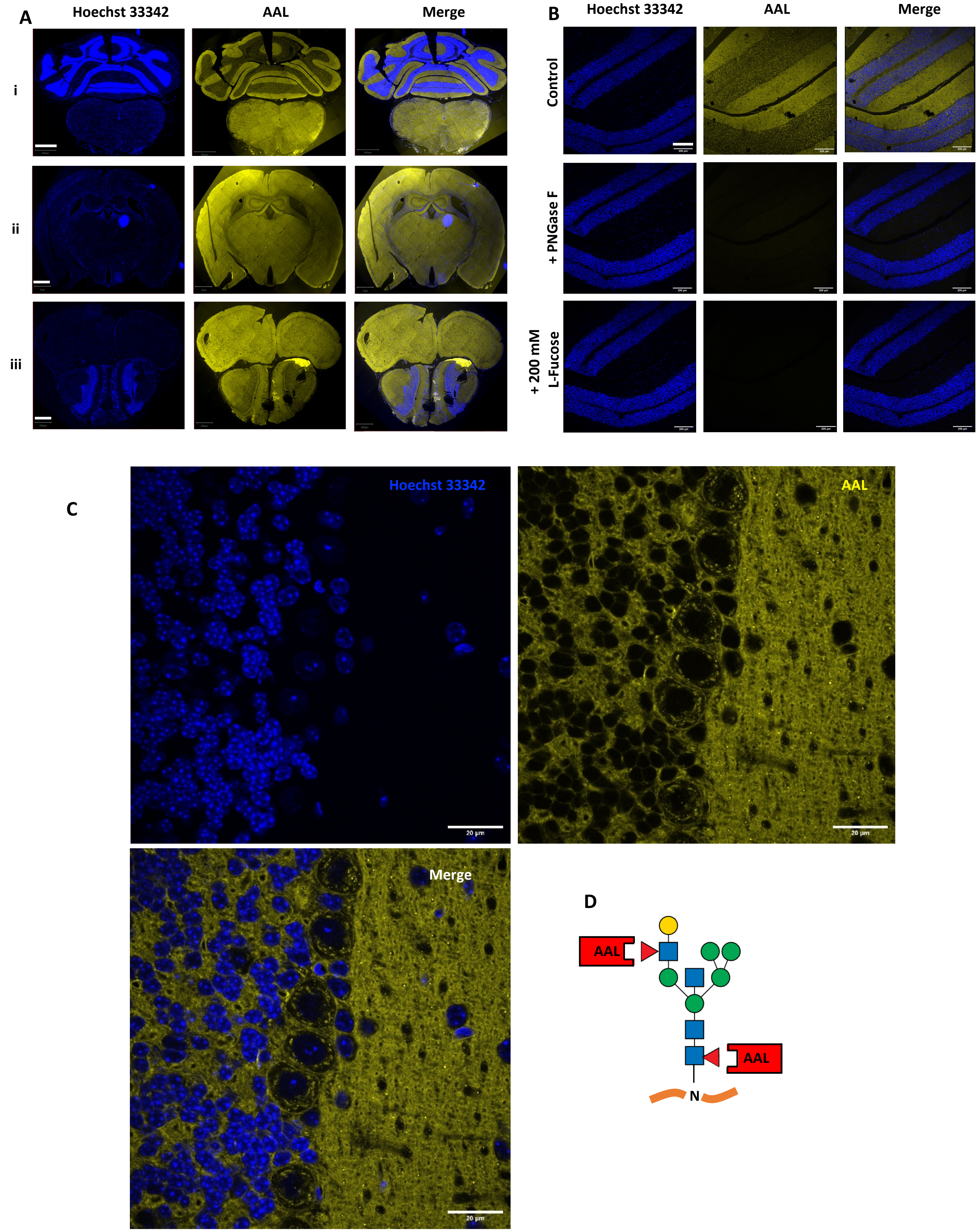


**Supplementary Figure 4 - AAL binding in the brain is broad with some regional differences. A)** AAL binding across multiple brain regions corresponding to coronal sections (i, ii, iii) indicated in Supp. Fig. 1. Scale bars i = 800 µm, ii = 1 mm, iii = 500 µm. **B)** AAL binding specificity in the cerebellum is confirmed by sensitivity to PNGase F and competition with L-fucose. Scale bar = 200 µm. **C)** AAL binding of the Purkinje cell layer, showing increased signal in the molecular layer. Scale bar = 20 µm. **D)** Binding preference of AAL to both core and antennary fucose of N-glycans.

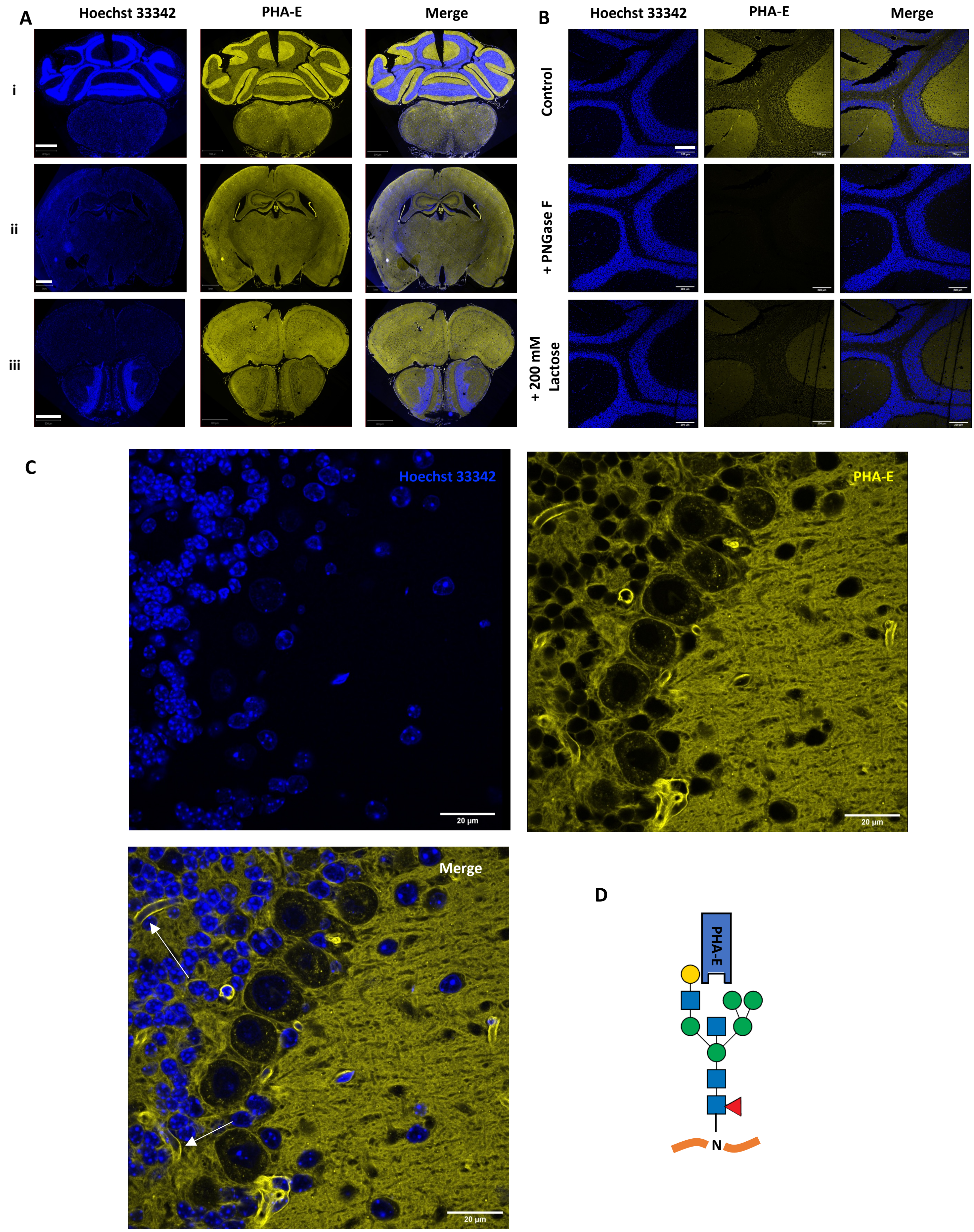


**Supplementary Figure 5 - PHA-E binding in the brain is broad with some regional differences. A)** PHA-E binding across multiple brain regions corresponding to coronal sections (i, ii, iii) indicated in Supp. Fig. 1. Scale bars i = 800 µm, ii = 1 mm, iii = 800 µm. **B)** PHA-E binding specificity in the cerebellum is confirmed by sensitivity to PNGase F and competition with lactose. Scale bar = 200 µM. **C)** PHA-E binding of the Purkinje cell layer, showing increased signal in the molecular layer and binding to luminal structures (white arrows). Scale bar = 20 µm. **D)** Binding preference of PHA-E to bisected N-glycans.

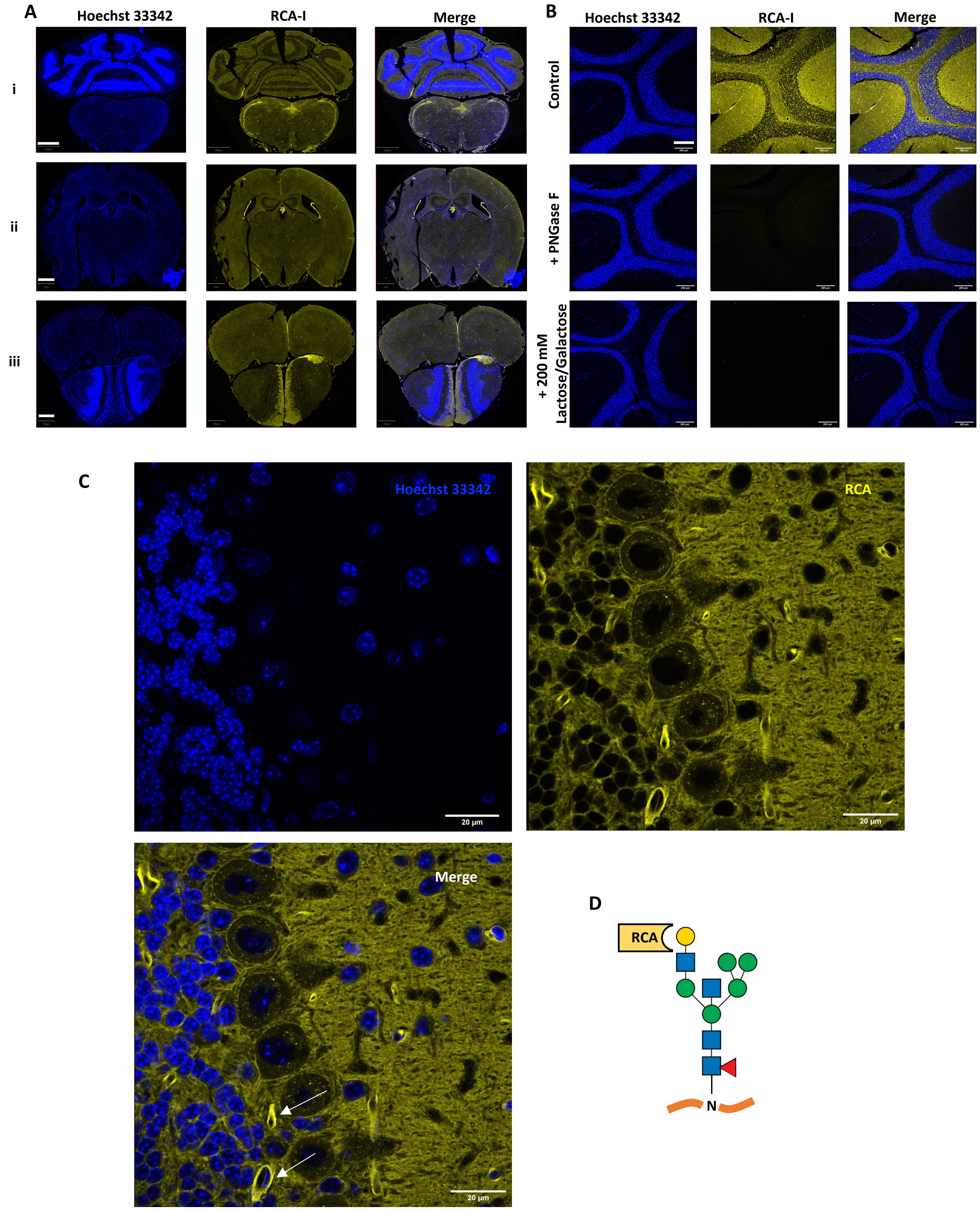


**Supplementary Figure 6 - RCA binding in the brain is scarce with increased binding to luminal structures. A)** RCA binding across multiple brain regions corresponding to coronal sections (i, ii, iii) indicated in Supp. Fig. 1. Scale bars i = 800 µm, ii = 1 mm, iii = 500 µm. **B)** RCA binding specificity in the cerebellum is confirmed by sensitivity to PNGase F and competition with lactose and galactose. Scale bar = 200 µm. **C)** RCA binding of the Purkinje cell layer, showing increased binding to luminal structures (white arrows). Scale bar = 20 µm. **D)** Binding preference of RCA to galactose of N-glycans.

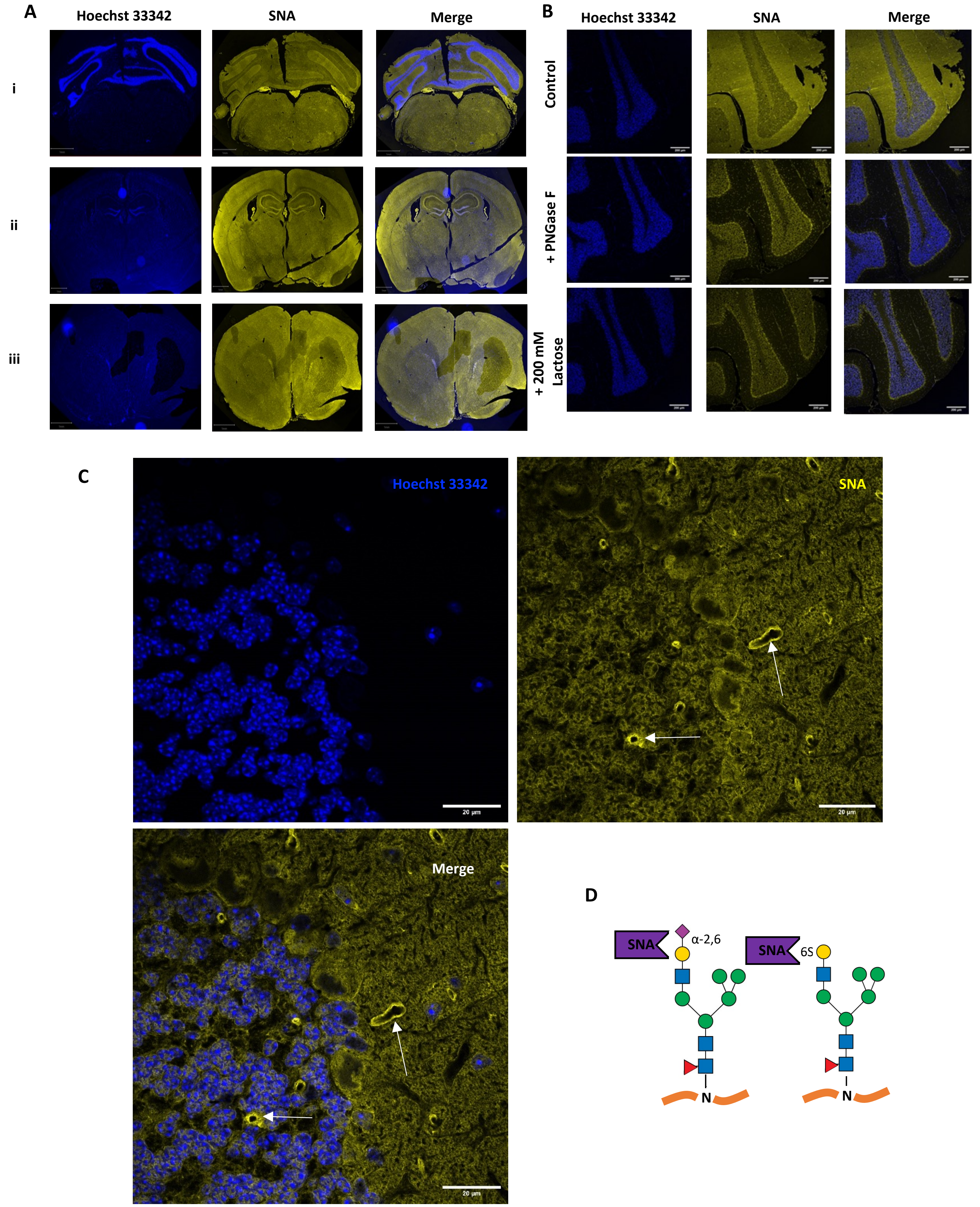


**Supplementary Figure 7 - SNA binding in the brain is diffuse with non-specific signal in the cerebellar granular layer. A)** SNA binding across multiple brain regions corresponding to coronal sections (i, ii, iii) indicated in Supp. Fig. 1. Scale bars i = 800 µm, ii = 1 mm, iii = 500 µm. **B)** SNA binding specificity in the molecular layer and arbor vitae of cerebellum is confirmed by sensitivity to PNGase F and competition with lactose, though non-specific binding in the granular layer is present in both conditions. Scale bar = 200 µm. **C)** SNA binding of the Purkinje cell layer, showing a patchy distribution with some signal detected in luminal structures (white arrows). Scale bar = 20 µm. **D)** Binding preference of SNA to N-glycans containing α-2,6 linked sialic acid. SNA may bind to the LacNAc structure with sialic acid based on previous reports and inhibition with lactose, as well as N-glycans containing 6-0-sulfated galactose.



.

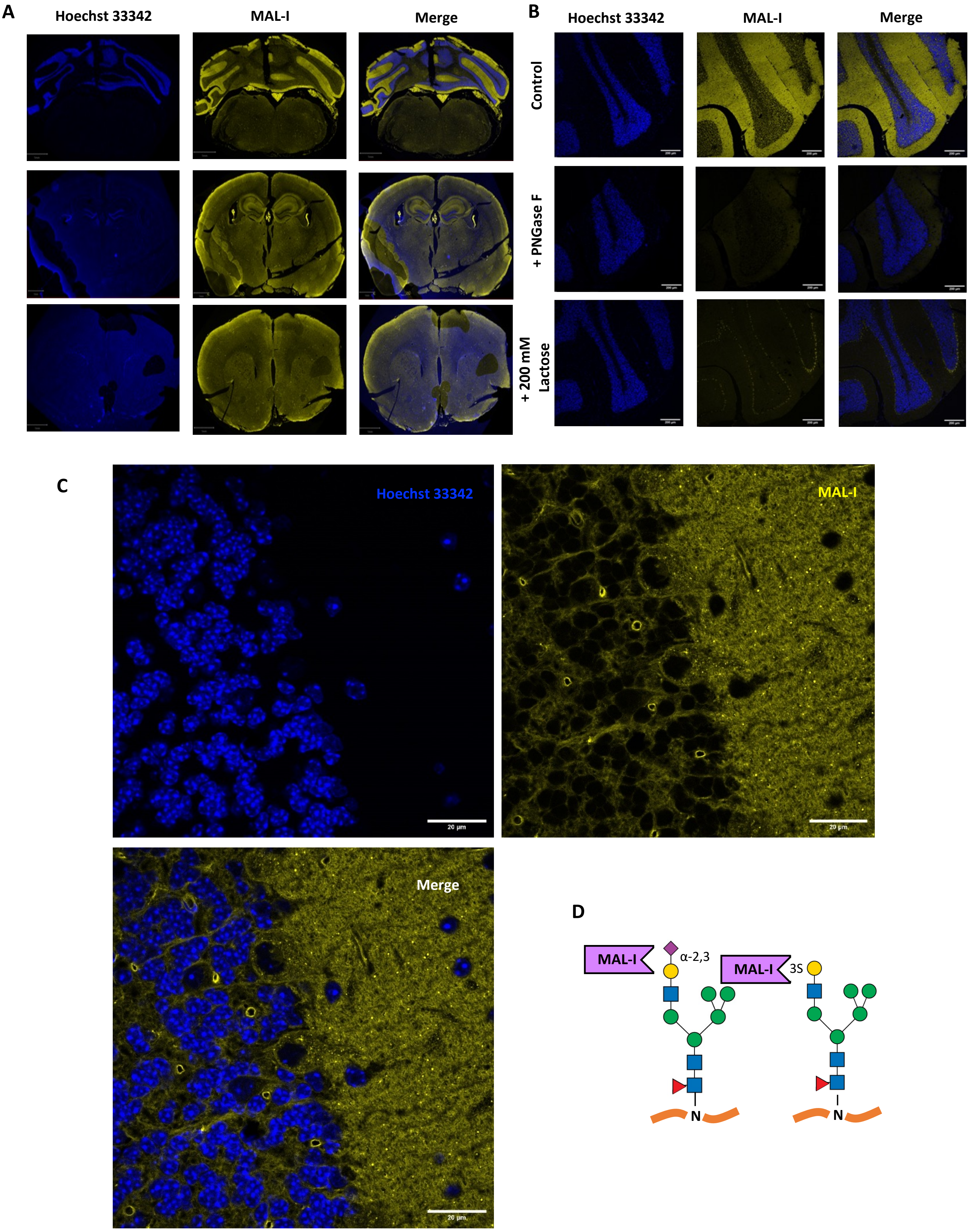


**Supplementary Figure 8 - MAL-I binding in the brain is enriched in the cerebellar molecular layer and the choroid plexus. A)** MAL-I binding across multiple brain regions corresponding to coronal sections (i, ii, iii) indicated in Supp. Fig. 1. Scale bars i = 800 µm, ii = 1 mm, iii = 500 µm. **B)** MAL-I binding specificity in the cerebellum is confirmed by sensitivity to PNGase F and competition with lactose. Scale bar = 200 µm. **C)** MAL-I binding of the Purkinje cell layer, showing increased binding to the molecular layer. Scale bar = 20 µM. **D)** Binding preference of MAL-I to N-glycans containing LacNAc with α-2,3 linked sialic acid, as well as N-glycans containing 3-O-sulfated galactose.
.
